# Comparative Structural Analyses of Selected Spike Protein-RBD Mutations in SARS-CoV-2 Lineages

**DOI:** 10.1101/2021.06.23.449639

**Authors:** Urmi Roy

**Author notes:** To whom correspondence should be addressed.; Phone: 315/212-7346.

## Abstract

The severity of the covid 19 has been observed throughout the world as severe acute respiratory syndrome coronavirus 2 (SARS-CoV-2) had spread globally claiming more than 2 million lives and left a devastating impact on peoples’ life. Recently several virulent mutant strains of this virus, such as the B.1.1.7, B.1.351, and P1 lineages have emerged. These strains are predominantly observed in UK, South Africa and Brazil. Another extremely pathogenic B.1.617 lineage and its sub-lineages, first detected in India, are now affecting some countries at notably stronger spread-rates. This paper computationally examines the time-based structures of B.1.1.7, B.1.351, P1 lineages with selected spike protein mutations. Additionally, the mutations in the more recently found B.1.617 lineage and some of its sub-lineages are explored, and the implications for multiple point mutations of the spike protein’s receptor-binding domain (RBD) are described.

## 1. Introduction

COVID 19, the severe acute respiratory syndrome coronavirus 2 (SARS-CoV-2) has claimed more than 2 million lives and left a devastating global impact since March 2020 [1]. In recent months, several variants of this life-threatening virus have been emerged with greater spread-rates, adaptability and fitness. Among these virulent lineages B.1.1.7, B.1.351 and P1 were initially detected in UK, South Africa and Brazil, respectively [2]. These nomenclatures are based on Pango lineage [3]. As summarized in Table 1, these “Variant of Concern” have several notable mutations, some of which are common among the three lineages The Table also includes the highly pathogenic B.1.617 lineage and some of its sub-lineages, that have been first found in India more recently, and already included in CDC’s “Variants of Interest” [4]. The delta variant (also known as B.1.617.2) has been listed under CDC’s “Variant of Concern” on June 15th, 2021. The present work focuses on a set of comparative structural analyses of these new SARS-CoV-2 variants.

**Table 1.** Selected Mutations in SARS-CoV-2 Lineages.

| Name of the lineage/sub-lineage | Spike protein mutations | Experiments with key mutations on SARS-CoV-2 spike RBD |
| --- | --- | --- |
| B.1.1.7<sup>[2]</sup> | 69/70 deletion, 144Y deletion, E484K*, S494P*, N501Y, A570D, D614G, P681H | E484K, S494P, N501Y |
| B.1.351<sup>[2]</sup> | K417N, E484K, N501Y, D614G | K417N, E484K, N501Y |
| P1<sup>[2]</sup> | K417T**, E484K, N501Y, D614G | K417T, E484K, N501Y |
| B.1.617<sup>[4]</sup> | L452R, E484Q, D614G | L452R, E484Q |
| B.1.617.1<sup>[4]</sup> | T95I*, G142D, E154K, L452R, E484Q, D614G, P681R, Q1071H |  |
| B.1.617.3<sup>[4]</sup> | T19R, G142D, L452R, E484Q, D614G, P681R, D950N |  |
| variant with possible mutations<sup>[16]</sup> | L452R, Y453F | L452R, Y453F |
| * found in some sequences<br>** Initially observed as K417N/T, later this mutation was identified as K417T in the P1 lineage |  |  |

Computational structural biology is rapidly becoming an integral part of applied immunology, as this field continues to aid the understanding of the structural basis of proteins, and thus, plays a key role in the development of preventive drug designs [5, 6]. Since the beginning of the Covid-19 pandemic, the availability of experimental results about the structure/function, epidemiological distribution and mutational fitness of this novel pathogen has been very limited in the commonly available literature. As a result, scientists have heavily relied on simulation-based tools and strategies to investigate this virus. In this regard, computational tools of immunoinformatics can be particularly useful to investigate such evolving infectious pathogens and host-pathogen interactions [7–9]. Our present effort is guided by these considerations.

In our previous papers, we have analyzed several biologically relevant protein structures and mutant models of the angiotensin peptide coordinated to the Zn-bound ACE2 receptor; more recently, we have reported a model structure of the SARS-CoV-2 N501Y variant [10–15]. In this current investigation of SARS-CoV-2 lineages, we will examine the implications for multiple point mutations on their spike RBD. In particular, we will measure the structural and conformational variations of these mutant variants as functions of time and, demonstrate how structural change corresponds to their functions.

The SARS-CoV-2 genome contains non-structural (NSPs) as well as structural proteins. There are 16 NSPs in SARS-CoV-2 genome, NSP1-NSP16. The structural portions consist of spike (S), envelope (E), membrane (M) and nucleocapsid (N) proteins. The S glycoprotein is the main interacting site for the host entry, and plays an important role in host defense and antibody neutralization. The accessory proteins contain several open reading frames (ORFs) including ORF1a/1b, ORF3a/3b, ORF6, ORF7a/7b, and ORFs8-10. The SARS-CoV-2 genome contains a single strand sense RNA, containing some similarities with the previously identified beta-coronavirus family members of the SARS-CoV and the Middle East respiratory syndrome (MERS-CoV) viruses.

Structural illustrations of SARS-CoV-2 and its genome structure are schematically presented in Fig 1a-b. Though it has some structural similarities with SARS and MERS, as observed in their sequence similarities, the fast transmissibility and adaptability of the highly pathogenic SARS-CoV-2 is rather unique. The SARS-CoV-2 is ~1273 amino acids (AAs) long, that contains the S1 (~14–685AAs) and S2 (~686–1273AAs) subunits. At the start of this structure, a small N-terminus signaling peptide (~1–13AAs) is present. Then comes the S1 subunit, that helps in receptor binding and comprised of two domains, N-terminal domain (NTD; ~14-305AAs) and receptor binding domain (RBD; ~319-541AAs) [17]. The mutations we discussed in this report are centered on the lately found variants of S1 RBD.

**Figure 1.**
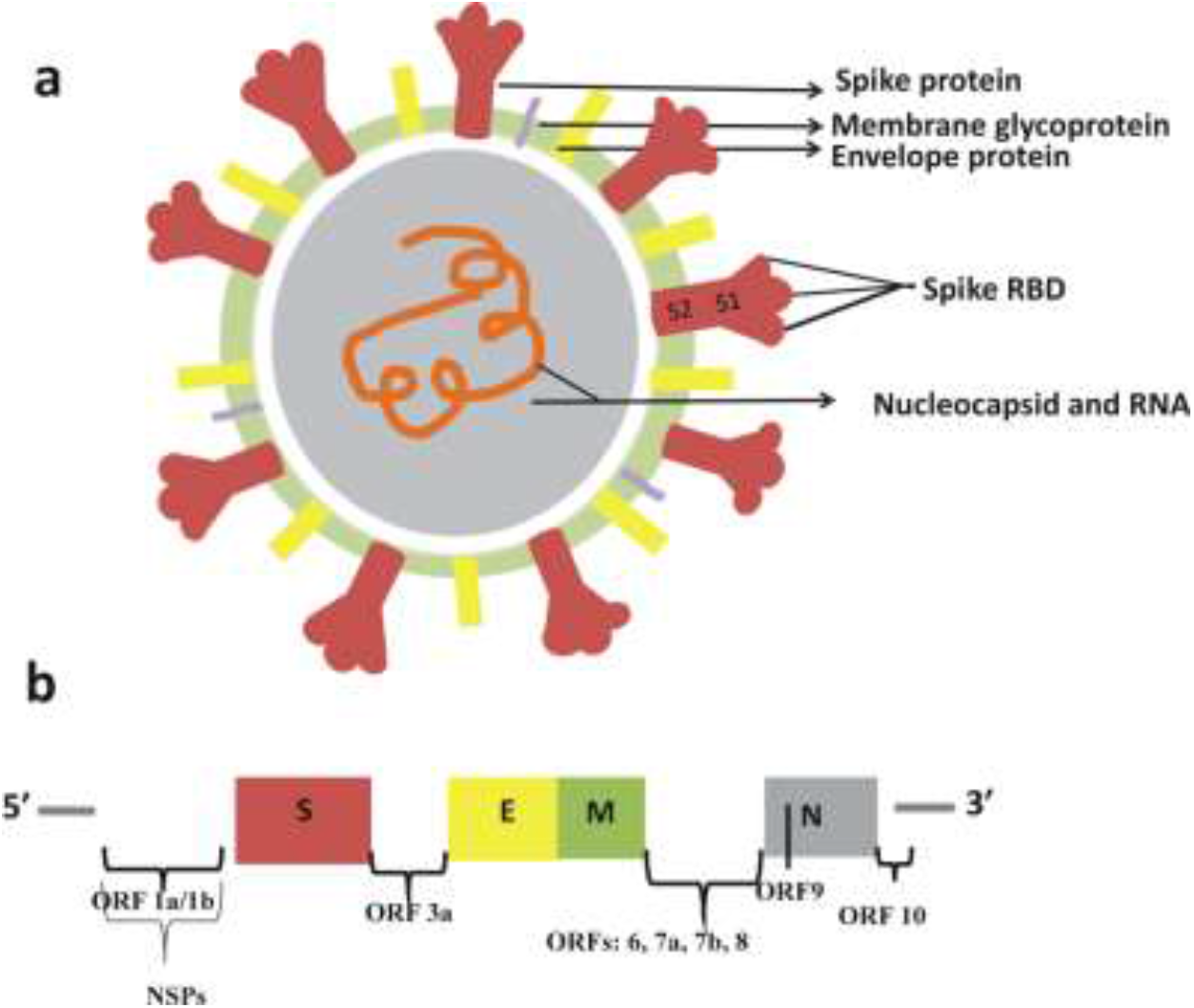
a. Schematic diagram of the SARS-COV-2 structure where the spike proteins are displayed in light brown. b. The genome structure of the SARS-COV-2.

## 2. Materials and Methods

We have used the spike protein RBD for the first set of simulations, where we have selected the wt type 6M0J: E as SARS-CoV-2 S1 RBD [18] and some of the reported mutations found in the B.1.1.7, B.1.351, P1 lineages. Additionally, we have analyzed the B.1.617, lineage, as well as the sub-lineages B.1.617.1/B.1.617.3 with selected common mutations in their S1-RBD. The B.1.1.7, B.1.351 and P1 strains are currently listed under CDC’s “Variant of Concern”, while B.1.617 and some of its sub-lineages are included in CDC’s “Variant of Interest”. Very recently, B.1.617.2, the delta variant has been listed under CDC’s “Variant of Concern”. A possible mutant with two single-point mutations in its S protein’s RBD is also examined [16].

Starting from the native structures (6M0J:E), the mutant variants were generated using the mutator gui of Visual Molecular Dynamics (VMD) [19]. In total we have simulations for the mutations listed in Table 1, one wt S1 RBD and five S1-RBD variants with selected mutations. These simulations have used the Nanoscale Molecular Dynamics (NAMD), quickMD and Visual Molecular Dynamics (VMD) software programs [19–21]. After completing the initial protocols (minimization/annealing and equilibration processes), the MD simulation was continued for 30 ns. The integration time was 2 fs for all procedures. All protocols have used the Generalized Born solvent-accessible surface area implicit solvation model [22]. For annealing and equilibration, the backbones were restrained, but no atoms were restrained during the final simulation process. Using Langevin dynamics the temperature was maintained at 300 K during final simulation process. The details of the simulation protocols are described elsewhere [15]. The proteins’ 3D models were set up by using Biovia’s Discovery Studio Visualizer [23].

## 3. Results and Discussion

Fig.1a shows a generalized schematic of the wt SARS-CoV-2 structure, where the S, E, M, N and the viral RNA, as well as the two subunits S1 and S2 of the S protein are identified. The mutations considered in this report are found in the RBD, within the S1 subunit of the S protein. Fig. 1b shows a typical genomic display of the SARS-CoV-2, where different nonstructural and structural parts are presented, along with the open reading frames (ORFs).

As shown in Fig 2 the structural illustrations of the wt and mutant variants of the SARS-CoV-2 RBD corresponds to the 6M0J:E subunit. This is a relatively small (319-541) subunit, and in Fig. 2a, we have focused on three selected mutant residues E484K, S494P and N501Y of the B.1.1.7 lineage. The mutant variant with residues K417N, E484K, N501Y of B.1.351 lineage is displayed in Figure 2b. The mutant variant with residues K417T, E484K, N501Y of P1 lineage is shown in Figure 2c. The two mutant residues L452R and E484Q of the B.1.617 lineage are depicted in Figure 2d. The last two mutations are common to the B.1.617.1 and B.1.617.3 sub-lineages. Another possible variant with two mutations L452R, Y453F, on the spike RBD is displayed in Figure 2e. From Figure 2 it is evident that some of these mutations are located on the proteins’ surface regions.

**Figure 2.**
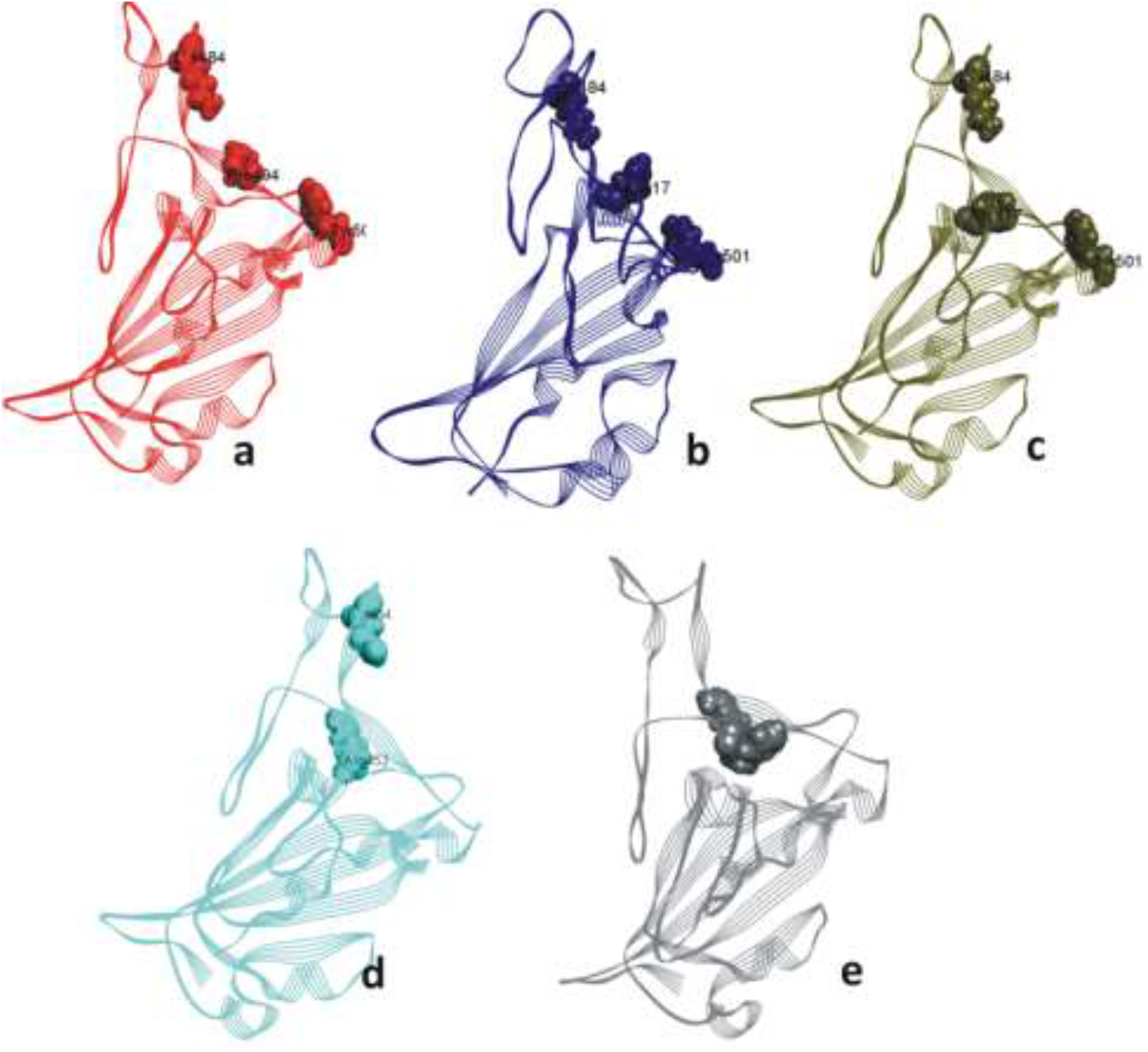
Secondary structures of SARS-CoV-2: S1-RBD variants with selected mutations. a. The selected mutant residues E484K, S494P and N501Y of B.1.1.7 lineage. b. The selected mutant residues K417N, E484K, N501Y of B.1.351 lineage. c. The selected mutant residues K417T, E484K, N501Y of P1 lineage. d. The mutant residues L452R, E484Q of B.1.617 lineage. e. The mutations L452R, Y453F within a possible mutant structure.

Figure 3a shows comparative RMSD plots of the SARS-CoV-2 variants with selected mutations based on the RBD of 6M0J. All these variants reached convergence, during the last 5 ns of the simulation. Nevertheless, as observed in the inset figure, the selected mutations with B.1.351 lineage (c) shows measurably higher RMSD values than those of P1 (d), B.1.617 (e) and the species of combined mutations 452 and 453 (f) during the last phase of the simulation. These last three variants are more stable than their WT species (a). Figure 3b represents RMSD of the selected mutations within the variants displayed in Figure 3a, and once again, the selected mutations of P1, B.1.617 and the third variant are seen to exhibit the lowest RMSD values, indicating a fairly stable nature shared by these mutations. The foregoing plots demonstrate that the selected mutations within P1, B.1.617 (also B.1.617.1 and B.1.617.3) lineages and the possible variant with 452 and 453 mutations are more stable than the mutant variants of B.1.351 and B.1.1.7. It is unknown, however, if the protein’s stability is dictated by the number of mutations within the variant. As in the case of the RMSD plots, the RMSF plots in Figure 3c also shows that, the B.1.617 lineage (e), and the L452R-Y453F mutations (f) are characterized by minimal fluctuations (lowest in their comparison group). At the same time, the selected mutations in B.1.351 lineage (c) once again show the higher range of fluctuations in Fig. 3c. We have plotted the RMSD and RMSF graph for the wt RBD structure previously but for comparison we have included it in Figure 3a and c [15]. Figure 3d shows the hydrogen bond numbers during the simulation time; for none of the cases considered, these numbers exhibit any significant variations.

**Figure 3.**
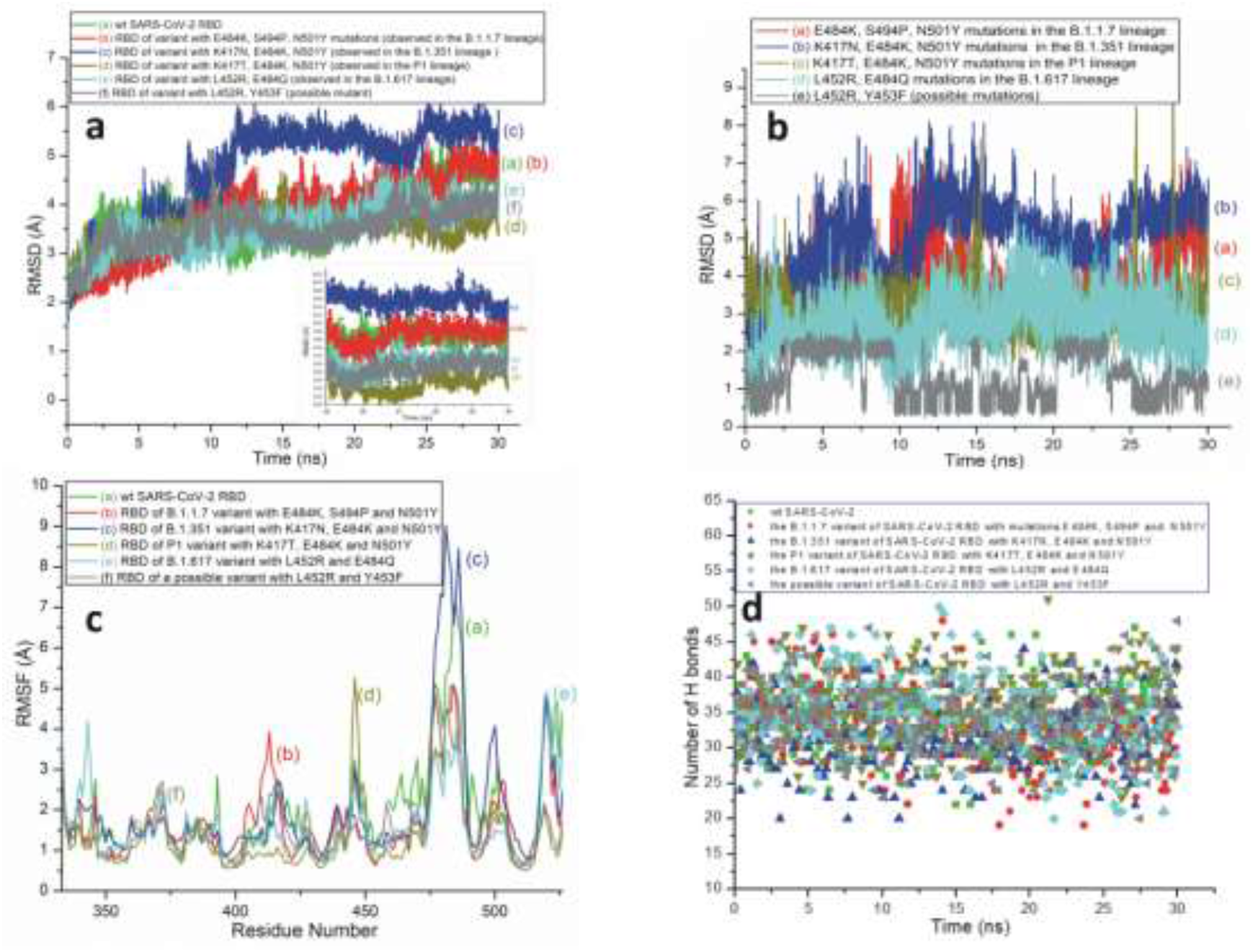
a. RMSD plots of wt and different SARS-CoV-2 variants with selected S1-RBD mutations. The two mutations identified in B.1.617 lineage (e) are also common in B.1.617.1 and B.1.617.3 sub-lineages. Inset showing the RMSDs for the last 5ns. b. The RMSD graph with selected mutant residues from different lineages. c. The RMSF plots for the wt and different SARS-CoV-2 variants with selected S1-RBD mutations. d. Variations in the number of H bonds of wt and different SARS-CoV-2 variants with selected mutations.

Figures 4-5 represent time based secondary structure-changes of the variant proteins as well as those of the selected mutant residues within these mutant strains. Secondary structures, in particular, the α helices and β sheets play a crucial role in determining proteins stability. From Figure, 4a-a’ it is clear that the mutations, S494P in B.1.1.7, is quite stable. Near the end of the simulation time, the E484K and N501Y transformations show slight fluctuations from coils to 3_10_ helices though they disappeared near the end the simulation. However, the actual manifestation of these latter effects may change with time.

**Figure 4.**
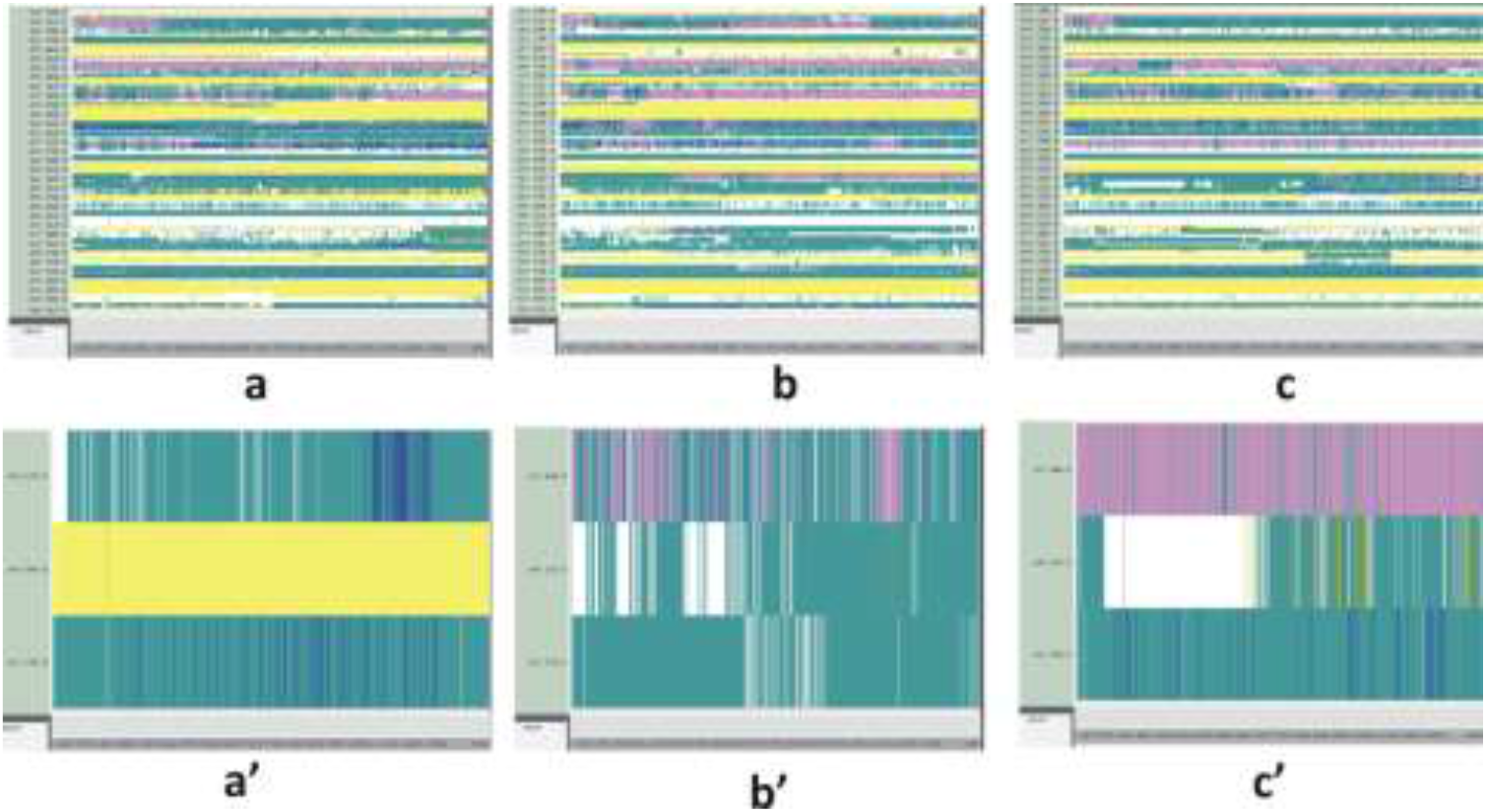
Time based secondary structure changes of SARS-CoV-2 mutant S1-RBDs. The time-based structure changes of a. the variant of SARS-CoV-2 RBD with mutations E484K, S494P and N501Y. a.’ E484K, S494P and N501Y residues within the B.1.1.7 lineage. b. the variant of SARS-COV-2 RBD with mutations K417N, E484K, N501Y. b’. The K417N, E484K, N501Y residues within the B.1.351 lineage. c. the variant of SARS-COV-2 RBD with mutations K417T, E484K, N501Y. c’. K417T, E484K, N501Y residues within the P1 lineage.

**Figure 5.**
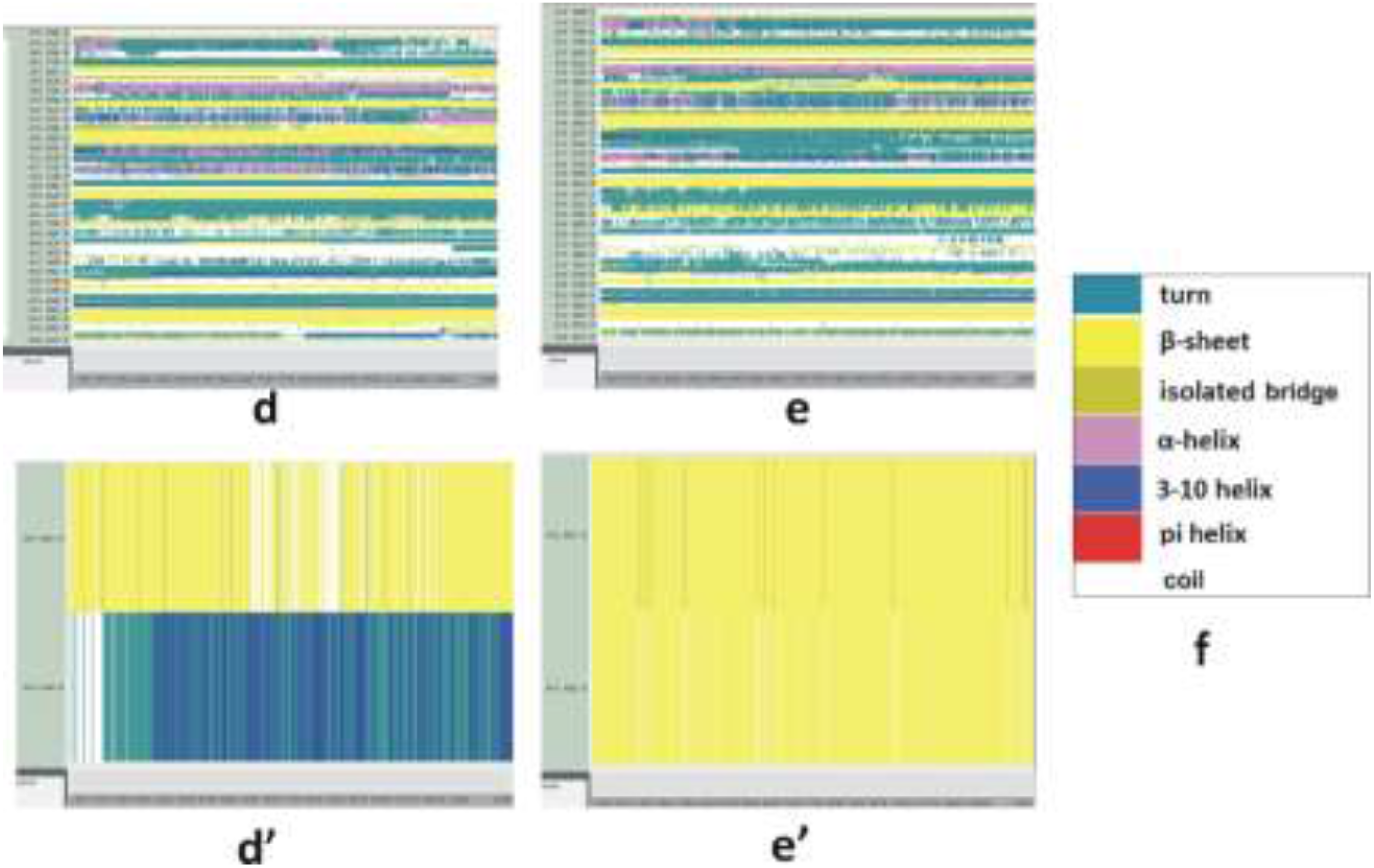
Secondary structure changes of d. the variant of SARS-COV-2 RBD with mutations L452R and E484Q. d’ L452R and E484Q residues within the B.1.617/B.1.617.1/B.1.617.3 lineage. e. the variant of SARS-COV-2 RBD with mutations L452R and Y453F. e’ L452R and Y453F residues within a possible mutant variant. f. The color code of secondary structure analyses.

Within the B.1.351 variant in Figure 4b-b’, the K417N mutation shows some variations from α helices to 3_10_ helices, turns and coils. This makes the K417 unstable during the last phase with the extinction of the α helices. The E484K mutation in this variant may show higher level of stability as some coils and turns are converted to few β sheets near the last phase of the simulation – even though they eventually disappear at the end. Coils to turns are still observed here at the end of simulation. There are some changes from turns to coils in the 501 residue, but it is not clear from the data if the latter correspond to the so called “coiled coils”.

Among the mutations we studied in P1, the K417T and N501Y do not show any significant structure variations throughout the simulation time. The E484K mutation exhibits a higher level of stability as some coils and turns in this species are converted to β sheets during the last phase of the simulation (Figure 4c-c’).

Within the L452R residue of B.1.617 lineage and the B.1.617.1/B.1.617.3 sub-lineages the intermittent 3-10 helices and the isolated bridges are completely transferred into β sheets making this mutant residue more stable. In the case of E484Q, the random coils and turns are converted to 3_10_ helices that are more stable, which, consequently make the secondary structure more rigid and solid (Figure 5d-d’).

The L452R and Y453F mutations in Figure 5e-e’ are stable and do not indicate any significant secondary structure changes during the simulation. The secondary structure change of a protein ligand is a key factor for fully understanding the protein’s functional changes. From Figures 4-5 it is evident that the selected mutant residues within the B.1.351 strain are fairly unstable, and that the residues within the B.1.617/B.1.617.1/B.1.617.3 variants are mostly stable. Among these mutations, L452R, Y453F, E484Q, S494P are quite stable, and E484K is particularly stable in the P1 strain.

Although numerous studies have already been reported on the structure and functions of SARS-CoV-2, the structures of the more recently found lineages have not been thoroughly investigated yet. The S1-RBD mutations of the B.1.617 lineage and its associated sub-lineages are of particular interest in this context. The B.1.617 strain was initially labeled as “double mutant” due to the presence of two mutations (E484Q, L452R) from two different lineages; L452R is found in B.1.427/B.1.429 while E484K, exists in both B.1.351 and P1. However, in B.1.617 the E484K mutation has been changed to E484Q. Another possible mutant variant with mutations L452R and Y453F are also the combination of two lineages, B.1.427/ B.1.429 and B.1.298.

The stable mutation in residue 452 may form a stronger complex with ACE2. The Y453F in the Figure 5e’ also exhibits stronger stability with time. The wt E484 residue has been recognized as a “repulsive residue” between the RBD-ACE2 complex [24]. Since the mutations in the E484K/Q residue are particularly stable in the P1 and B.1.617 strains (Figure 4c’and 4d’), the mutations in this residue may form a stronger bond with the receptor. S494P is also very strong as it resides within the β helix; within the B.1.1.7 lineage this appears to be the most stable residue. There are numerous examples in the literature that N501Y mutation forms a stable connection with the receptor.

## 4. Conclusions

According to the results presented here, the mutant RBD variant of B.1.617 (as well as some of its sub-lineages), P1 and the potential variant with two possible mutants are the most stable forms. Among the mutations we have studied in this work, L452R, Y453F, E484Q, S494P are fairly stable. N501Y does not show significant variations during the simulation timescale. The E484K within the P1 strain is also rather stable. Since these newly found lineages are more spreadable than their predecessor species, some of the mutations may escape from antibody neutralization and cellular immunity. In fact, some of the variants with mutations K417N/T, E484K, L452R and Y453F are recognized as antibody neutralizing escape mutants [16, 25]. The steady mutations identified here to occur within the highly infective species may help to further understand for the associated antibody cross-reactivity, and may also facilitate the task of designing effective inhibitors. The enhanced stabilities of some of the mutant residues, as found here with these newer variants, may have implications in the context of future vaccine developments to combat other impending strains and pathogenic variants of SARS-CoV-2.

## Acknowledgments

The author acknowledges utilization of the following simulation and visualization software packages: 1) NAMD and 2) VMD: NAMD and VMD, developed by the Theoretical and Computational Biophysics Group in the Beckman Institute for Advanced Science and Technology at the University of Illinois, Urbana-Champaign. 3) Discovery Studio Visualizer: Dassault Systèmes BIOVIA, Discovery Studio Modeling Environment, San Diego, CA: Dassault Systèmes (2015).

